# Amplitude of circadian rhythms becomes weaker in the north, but there is no cline in the period of rhythm in a beetle

**DOI:** 10.1101/2020.08.28.272070

**Authors:** Masato S. Abe, Kentarou Matsumura, Taishi Yoshii, Takahisa Miyatake

## Abstract

Many species show rhythmicity in activity, from the timing of flowering in plants to that of foraging behaviour in animals. The free-running periods and amplitude (sometimes called strength or power) of circadian rhythms are often used as indicators of biological clocks. Many reports have shown that these traits highly geographically variable, and interestingly, they often show latitudinal or altitudinal clines. In many cases, the higher the latitude is, the longer the free-running circadian period (i.e., period of rhythm) in insects and plants. However, reports of positive correlations between latitude or longitude and circadian rhythm traits, including free-running periods, the power of the rhythm and locomotor activity, are limited to certain taxonomic groups. Therefore, we collected a cosmopolitan stored-product pest species, the red flour beetle *Tribolium castaneum*, in various parts of Japan and examined its rhythm traits, including the power of the rhythm and period of the rhythm, which were calculated from locomotor activity. The analysis revealed that power was significantly lower for beetles collected in northern areas compared with southern areas in Japan. However, it is worth noting that the period of circadian rhythm did not show any clines; specifically, it did not vary among the sampling sites, despite the very large sample size (n = 1585). We discuss why these cline trends were observed in *T. castaneum*.

## Introduction

Latitudinal clines are of evolutionary interest because they indicate the action of natural selection [1]. Many traits correlate with latitude, for example, body size [2,3] and life history traits [4]. Circadian rhythm traits are also correlated with latitude.

Circadian rhythms are particularly important for timing or regulating key biological events in insects [5]. The free-running period and power of the rhythm (i.e., sometimes called the amplitude or strength of the rhythm) are often used as indicators of circadian rhythms [6,7], and there is much evidence that the free-running periods of circadian rhythms exhibit latitudinal or longitudinal clines at the phenotype to molecular levels in many taxonomic groups [8–12].

Additionally, in insect species other than *Drosophila*, many studies on the relationships between free-running periods and latitude have been conducted. In the linden bug, *Pyrrhocoris apterus*, higher-latitude populations are reported to have longer free-running periods [13]. Similar positive relations between latitude and circadian rhythms have been reported in a parasitic wasp, *Nasonia vitripennis* [12], and also in plant species [14].

In *Drosophila* species that evolved in tropical regions and then expanded their distribution to temperate regions, rhythm traits vary depending on where the flies live [15]. For example, pupal-adult eclosion rhythms in the far north were more arrhythmic than those in the south among *Drosophila littoralis* populations [16]. Circadian locomotor rhythms of *D. melanogaster* derived from Africa had a stronger power of rhythm than those of more northern *Drosophila* species in Europe [17]. However, pioneer studies revealed a negative relation between latitude and these traits [18]. Overall, these studies reveal that this topic remains controversial.

Given these discrepancies, we need more biological models. The red flour beetle *Tribolium castaneum* (Herbst) [Tenebrionidae] is a cosmopolitan stored-product pest [19], and hence, it can serve as an insect model species. Its biology and behaviour are well studied [20,21]. Furthermore, genome sequences are available for *Tribolium castaneum* [22].

In Japan, this species is distributed in most areas except Hokkaido, in the northern parts, meaning that it can be collected from a wide range of latitudes [23]. It provides us with a novel, fascinating model with which to examine the relations between latitude/longitude and circadian rhythm traits. Hence, we studied the relationships between latitude or longitude and circadian rhythm parameters, including the period of circadian rhythms, power of the rhythm, and locomotor activity, in *T. castaneum*.

## Materials and Methods

### Insects

The geographical populations of *Tribolium castaneum* used in the present study were collected from 38 different fields in Japan (Figure 1). Table S1 shows the latitude and longitude of the collection points, along with the number of samples. The northernmost and southernmost points where the insect could be collected were Aomori Prefecture (north latitude 40.89, east longitude 127.69) and Kumamoto Prefecture-C (north latitude 32.57, east longitude 130.66), respectively. The collection of insects was conducted during 2016 and 2017. We collected beetles from each rice pearling mill in Japan that was set up in a village or town with rice fields. More than twenty beetles were collected from each mill. The collected beetles were reared with a mixture of whole meal (Yoshikura Shokai, Tokyo) enriched with brewer’s yeast (Asahi Beer, Tokyo) and maintained at 25°C with a 16 h photoperiod (lights on at 07:00, lights off at 23:00). Each collected group was kept in a separate plastic Petri dish (90 mm in diameter, 15 mm in height). Since this species is a stored-grain pest, these laboratory conditions were chosen to closely mirror the native environment of these beetles. Before the experiments, each beetle population was reared for more than two generations in incubators (MIR-153, Sanyo, Osaka, Japan). We used beetles reared for a few generations in the incubator in the experiment.

**Figure 1.**
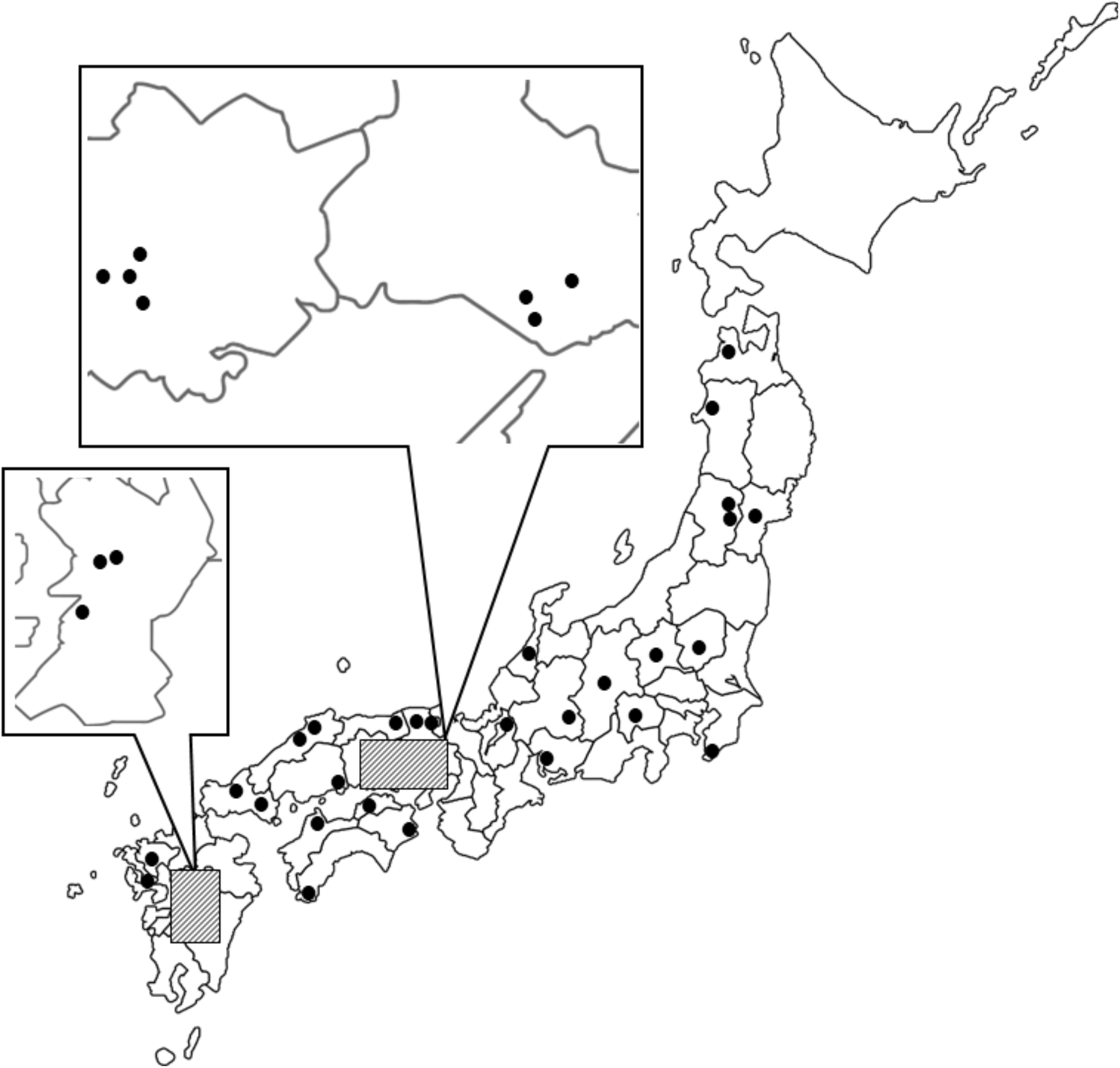
Collection locations of wild *T. castaneum* populations in Japan.

### Locomotor activity

To assess circadian rhythm, we maintained beetles under 16L:8D conditions for more than 20 days in an incubator kept at 25°C before the measurement of locomotor activity, and we then measured the locomotor activity of *T. castaneum* for 10 days in darkness. A beetle from each population was placed in a clear plastic Petri dish (30 × 10 mm) in an incubator (MIR-153, Sanyo, Osaka, Japan) maintained at 25°C under complete darkness (DD). The locomotor activity of each individual was monitored using an infrared actograph. An infrared light beam was passed through a clear Petri dish, and the beam was projected onto a photomicrosensor (E3S-AT11; Omron, Kyoto, Japan) that detected all interruptions of the light beam. Signals of interruption of the infrared light beam were recorded every 6 min [24]. The sample size of each population is shown in Table S1.

### Statistical analysis

To determine the circadian rhythm, the locomotor activity data collected for 10 days in constant dark conditions were analysed. The free-running period of circadian rhythms was established using a *χ^2^* periodogram test [25] for data on locomotor activity between 20 and 28 h [26]. Circadian rhythmicity was determined using *χ^2^* periodgram analysis, and “power” was used as an index of the strength of rhythms. The power of circadian rhythms was defined as the maximum difference between the *χ^2^* value and the significance threshold line at *P* = 0.05, that is, the size of the peak above the 5% threshold; see Figure 1 in [6]. Power is high when the rhythm is clear and strong, and a power of less than 0 indicates a statistically arrhythmic state. Moreover, total activity was calculated as the total number of interruptions of the infrared light over 10 days. To analyse the effects of temperature, latitude and sex on the period and power of the circadian rhythms and total activity, we used GLMs with Gaussian link functions. All statistical analyses were performed in R version 3.4.3 [27].

## Results

First, we analysed the relationship between the geographical area and the rhythm of power. To avoid multicollinearity between latitude and longitude (the correlation coefficient (*r*) between them was 0.83), we used one of the two as an explanatory variable.

The GLM results revealed a significant relationship between latitude and power (Table 1, Figure 2A), while sex was not significantly associated with power (Table 1). The estimated coefficient for latitude in the model was negative, suggesting that the higher the latitude was, the weaker the rhythm was. For longitude, we did not find a significant relationship (Table 1, Figure 2B).

**Figure 2.**
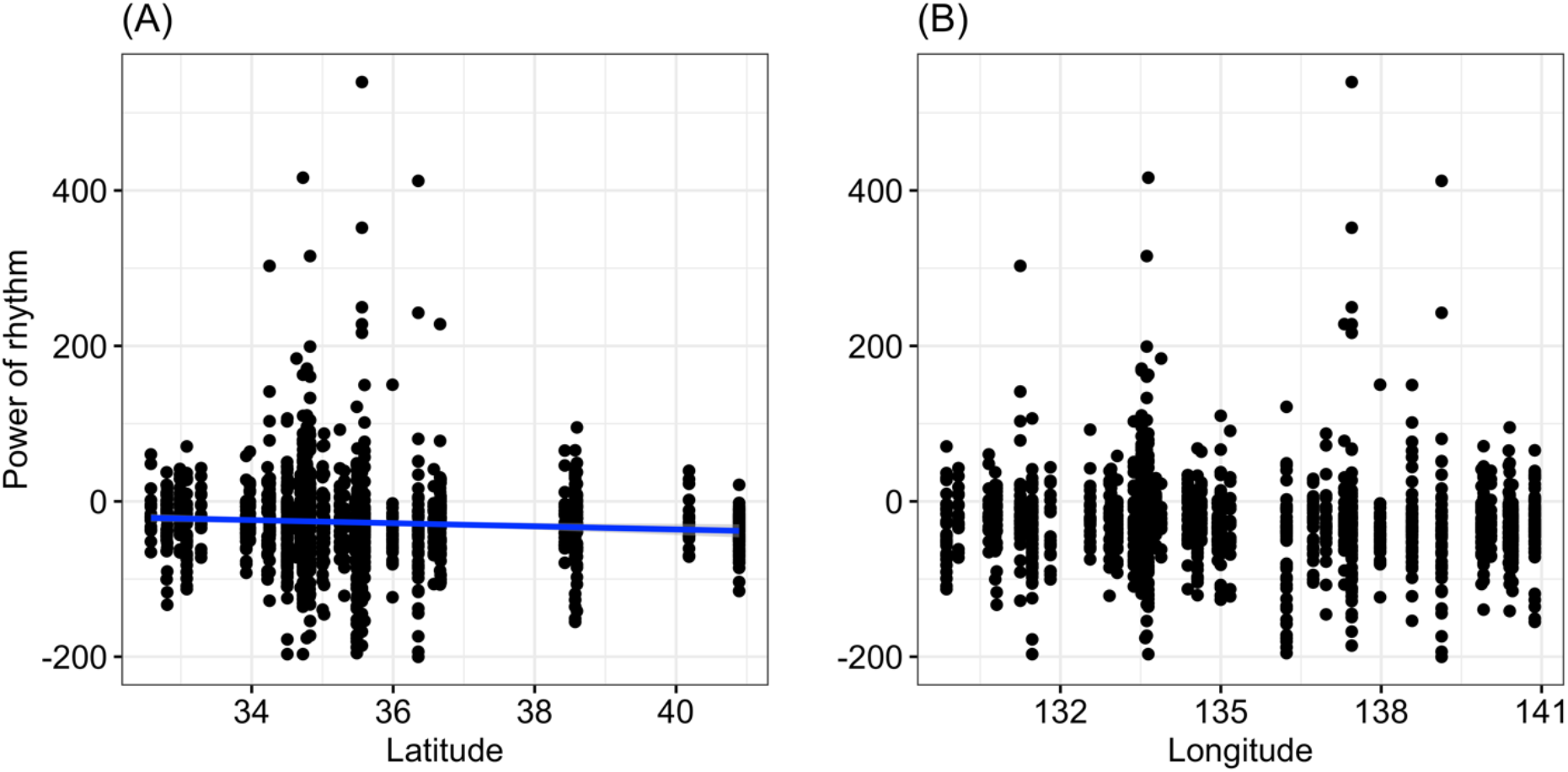
Relationship between the power of the rhythm and latitude (A) or longitude (B). The blue line represents the statistically significant regression line.

Second, we investigated the relationship between geographical area and the estimated period of circadian rhythms. The statistical results yielded no significant relationships between them (Table 1, Figure 3).

**Figure 3.**
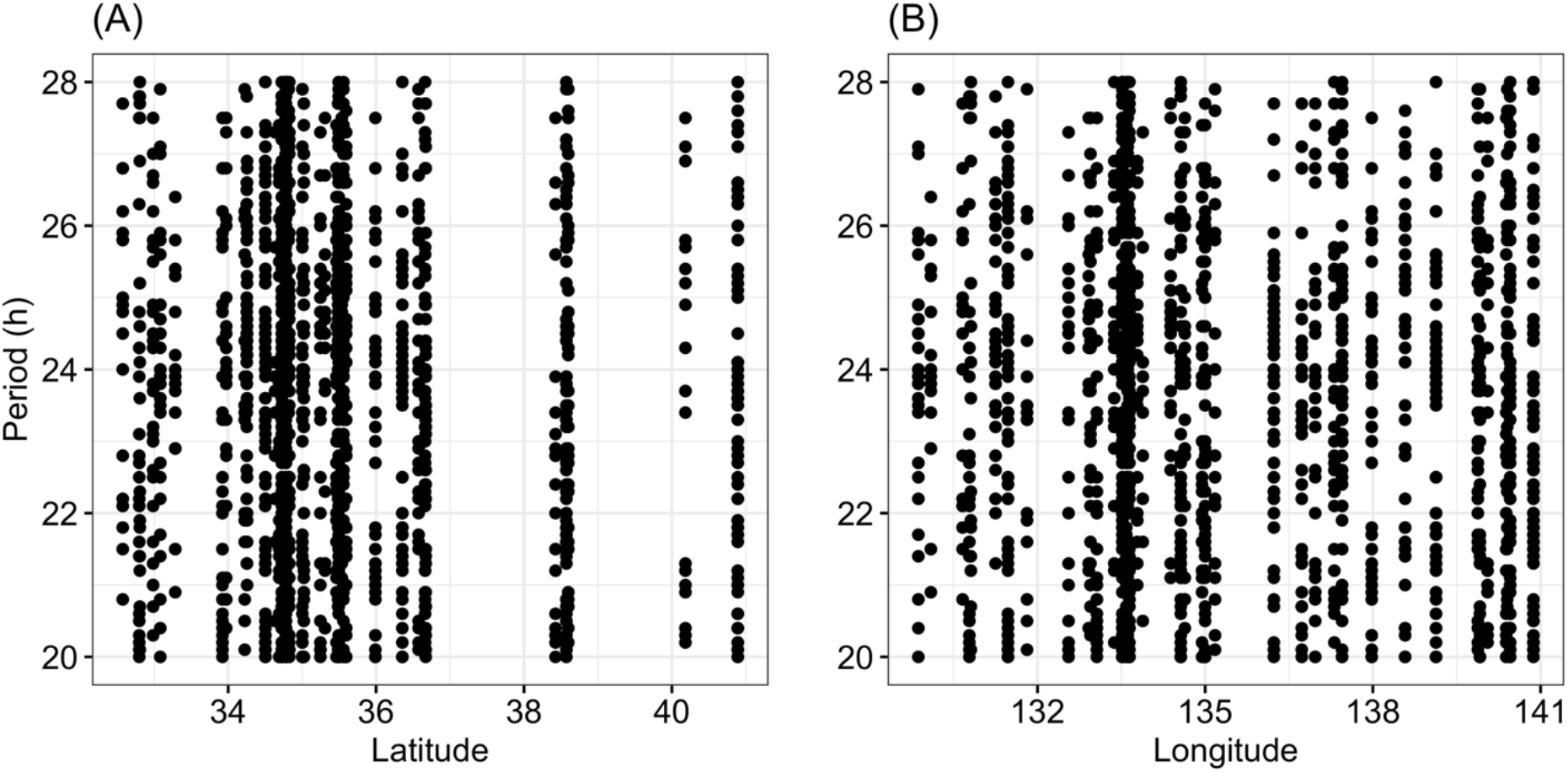
Relationship between the period of the rhythm and latitude (A) or longitude (B).

Finally, we investigated the relationship between geographical area and total activity (Figure 4). As the total activity had some outliers, we used log-transformed values. The results showed that total activity was associated with latitude and sex (Table 1, Figure 4B).

**Figure 4.**
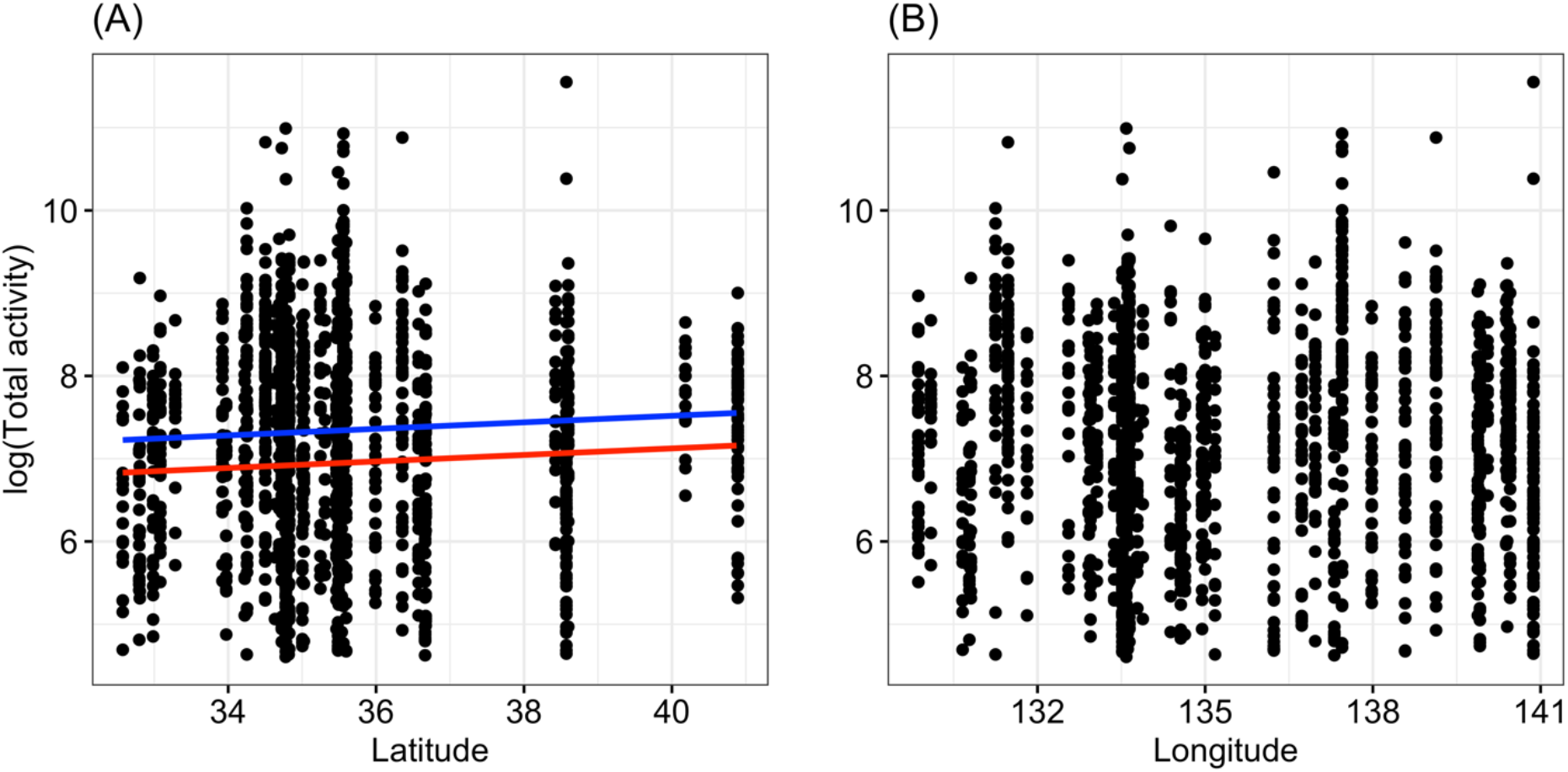
Relationship between total activity and latitude (A) or longitude (B). The blue and red lines represent the statistically significant regression lines of males and females, respectively.

## Discussion

In the present study, the period and power of circadian rhythms and locomotor rhythm varied among geographical populations of *T. castaneum*. Circadian periods seemed to vary evenly between 20 h and 28 h (Figure 3). Circadian rhythm variation occurs in various organisms [5]. The present results showed that the power of circadian rhythms was significantly lower for beetles collected in northern areas compared with southern areas (Figure 2). This result suggests that beetles collected from different parts of Japan have genetically different characteristics. In this study, we reared individuals collected from the fields for a few generations in a chamber under the same environmental conditions in the laboratory before measuring their traits. Therefore, it is unlikely that the present results were influenced by any maternal effects. On the other hand, no clines have been observed in the length of circadian rhythms.

The trend of weaker circadian rhythms in northern populations has also been observed in other insect species. Specifically, a clear rhythm was shown at lower latitudes whereas no rhythmic activity was shown at higher latitudes in *Hymenoptera* and *Drosophila* species [28,29]. Therefore, the present results are consistent with the results of these previous studies. Why is the rhythm weakened at higher latitudes? One answer may be that in more extreme environments, it may be easier to survive with less restriction of activity by the clock and more control by direct environmental responses, namely, masking of circadian activity [30,31]. To the best of our knowledge, clines in the power of the rhythm have been observed in only a few species.

On the other hand, positive relations between latitude and the length of circadian rhythms have often been reported; higher-latitude populations are reported to have longer free-running periods [12–14]. However, in the present study, no significant relationship was observed between the period of circadian rhythms and latitude or longitude, despite the very large sample sizes.

The cline in the amplitude (i.e., power) of the circadian rhythm in *T. castaneum* clearly suggests geographic variation. This result is very interesting considering the history of the controversy surrounding the dispersal distance of this insect, specially, that studies do not agree on its dispersal characteristics. Some studies suggest that this beetle disperses very well [32] suggested very high levels of active dispersal through adult flight in *T. castaneum* based on microsatellite genotypes. Drury et al. [33] reported that although these beetles have wings, dispersing by flying is rare at 25°C. On the other hand, Arnold et al. [34] suggested that this beetle usually travels by walking more often than it flies. Semeao et al. [35] showed that populations of *T. castaneum* collected from mills show spatial genetic structure, indicating the occurrence of a recent bottleneck in each mill. The present results clearly showed geographical variation in the amplitude of circadian rhythms among local populations in each mill.

We considered two few hypotheses regarding why no clines in the length of circadian rhythms were found in *T. castaneum*, as follow: bottlenecks and local adaptation. A small number of individuals or one fertilized female can enter and settle in individual rice mills scattered in the countryside of Japan [36]. A small number of *T. castaneum* will form each population within each mill. Predation pressures, including that from predator insects [36], and the differences due to human cleaning will differ among mills. These pressures can cause differences in traits, especially in the activity traits, of *T. castaneum*. Such selection pressures (local adaptation) and founder effects (bottlenecks) would cause a large degree of variation among *T. castaneum* populations. *T. castaneum* cannot fly under stable conditions. Indeed, as described above, although these beetles have wings, dispersing by flying is rare at 25°C [33], with walking being the most frequent mode of travel [34], and walking is the mechanism by which males locally search for females [37]. Additionally, Semeao et al. [35] showed that populations of *T. castaneum* collected from mills showed spatial genetic structure, indicating the occurrence of a recent bottleneck in each mill. Therefore, each mill used in the present study may be an ideal environment if without human cleaning, and thus, the beetles may not need to disperse from the rice bran in each mill.

On the other hand, adults of *T. castaneum* are known to fly well above 28°C [38]. There may be individuals flying beyond the area in the summer season in Japan. If gene flow is greater than expected, it might explain the lack of latitudinal and longitudinal clines shown in the present study. Ridley et al. [32] estimated the dispersal distance of *T. castaneum* using microsatellites and found that adults could fly at least 1 km per year in fields. They reported that *T. castaneum* is predominantly aggregates around areas of grain storage but actively disperses by flight between these spatially separated resources [32].

Another study showed that genetic distance was not significantly correlated with geographic distance among *T. castaneum* populations in mills in the United States [35]. Semeao et al. [35] provided evidence that populations of *T. castaneum* collected from mills showed spatial genetic structure, but the poor ability to assign individuals to source populations and the lack of isolation by distance suggested lower levels of gene flow than originally predicted. Konishi et al. [36] also suggested that different anti-predator strategies have evolved in each storehouse with and without predatory insects. These studies suggest that the dispute between gene flow or the possibility of evolution in individual storehouses in field mill systems will continue. The present result of a cline in amplitude suggests that gene flow is not occurring at the area scale that we examined. However, estimation of the degree of gene flow between rice mills and phylogenetic relationships between populations within the species in Japan is required.

Notably, the results of the present study showed significantly smaller *P*-values, even for small effect sizes, due to the very large sample size (*n* = 1585, total sample size). This suggests that the power of circadian rhythms is smaller at higher latitudes, but the change is not large. Moreover, it is worth noting that the period of circadian rhythms does not change, even with the very large sample size, in this beetle species. Further studies on circadian rhythms at the molecular level using this model beetle, which shows different phenotypic phenomena than other insect species, are needed in the future.

**Table 1.**
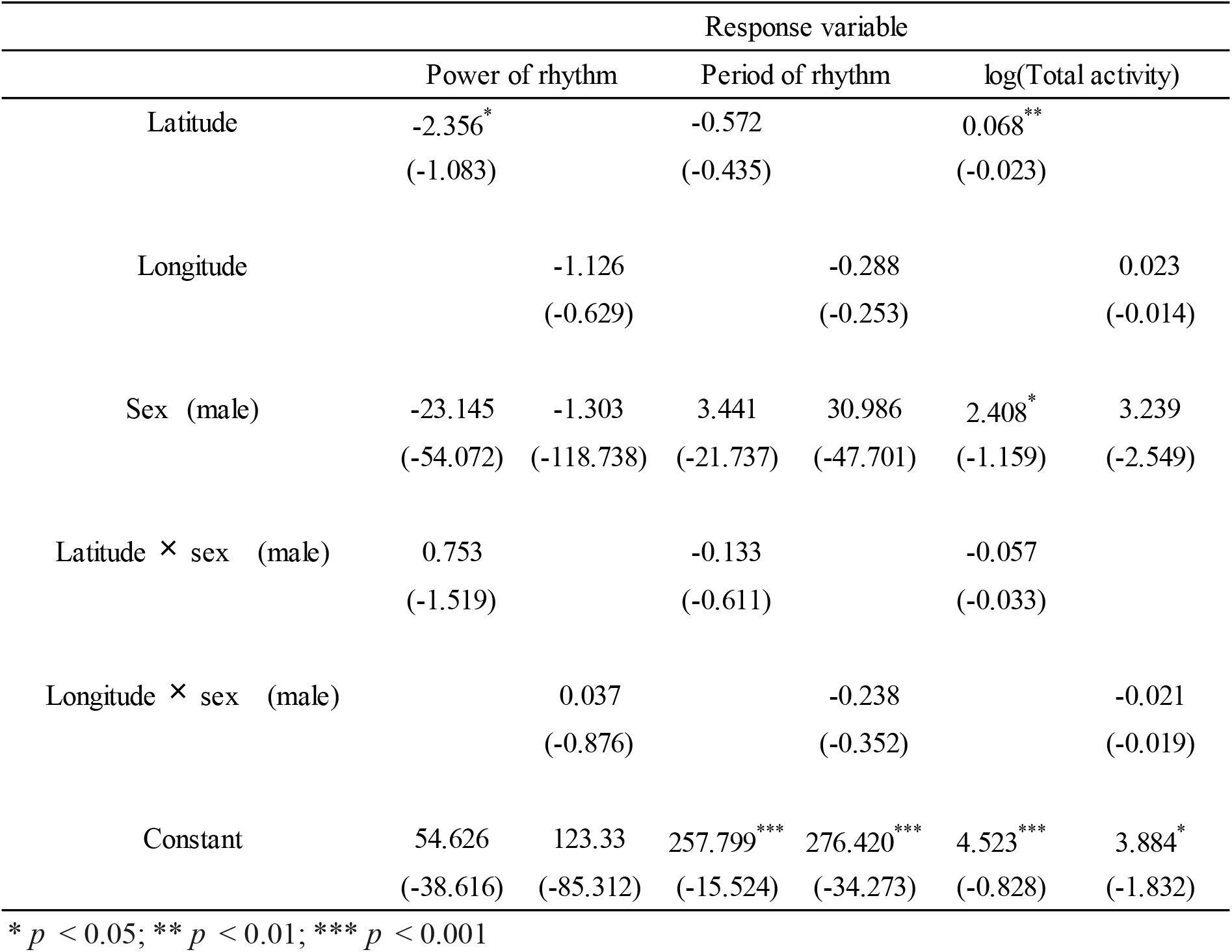
Statistical test results. Coefficients and standard errors obtained from GLM analysis are shown.

## Acknowledgement

We appreciate Yusuke Tsushima and Kouhei Nakao for insect sampling. This work was supported by the Japan Society for the Promotion of Science Grantin-Aid for Scientific Research Grants 16K14810 and 18H02510 to TM.

## Authors’ contributions

The experiments were designed by TM, and these were performed by KM. The data were analyzed by MSA. The manuscript was written by MSA, TY and TM. All authors approved the final version prior submission.

## Ethical Note

The populations of *T. castaneum* used in this study were collected from each mill located at 38 different fields in Japan. These populations have been maintained on wholemeal flour enriched with yeast at 25 °C under a 16:8 h light:dark cycle. These laboratory conditions closely resemble natural conditions of this stored product pest. All individuals in the experiment were handled with care and handling time was kept to an absolute minimum. The use of these beetles conforms to the Okayama University’s Animal Ethics Policy.

---

**Table S1.**
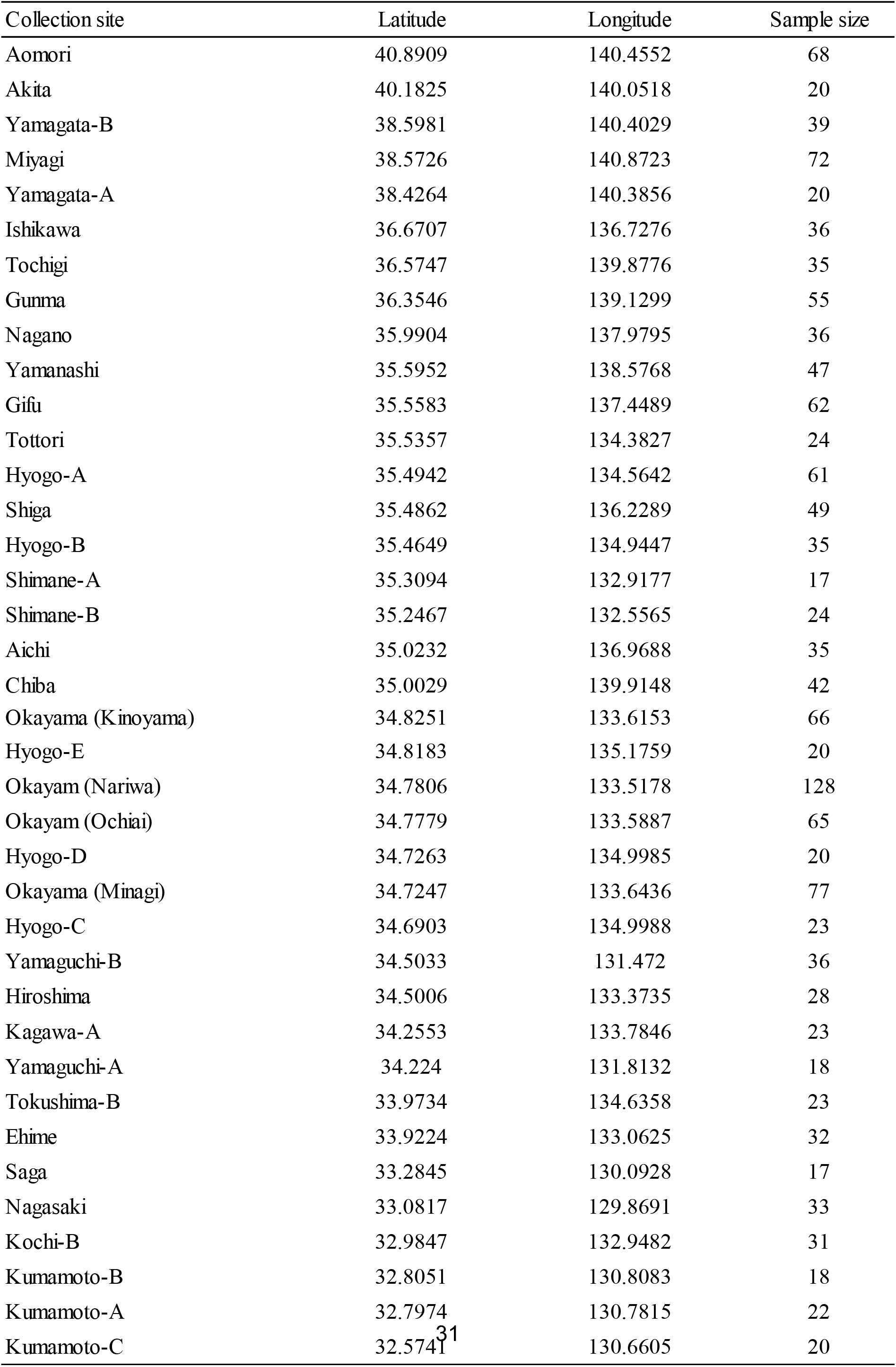
Collection site of *T. castaneum* populations with latitude and longitude.

